# RNA tertiary structure energetics predicted by an ensemble model of the RNA double helix

**DOI:** 10.1101/341107

**Authors:** Joseph D. Yesselman, Sarah K. Denny, Namita Bisaria, Daniel Herschlag, William J. Greenleaf, Rhiju Das

## Abstract

Over 50% of residues within functional structured RNAs are base-paired in Watson-Crick helices, but it is not fully understood how these helices’ geometric preferences and flexibility might influence RNA tertiary structure. Here, we show experimentally and computationally that the ensemble fluctuations of RNA helices substantially impact RNA tertiary structure stability. We updated a model for the conformational ensemble of the RNA helix using crystallographic structures of Watson-Crick base pair steps. To test this model, we made blind predictions of the thermodynamic stability of >1500 tertiary assemblies with differing helical sequences and compared calculations to independent measurements from a high-throughput experimental platform. The blind predictions accounted for thermodynamic effects from changing helix sequence and length with unexpectedly tight accuracies (RMSD of 0.34 and 0.77 kcal/mol, respectively). These comparisons lead to a detailed picture of how RNA base pair steps fluctuate within complex assemblies and suggest a new route toward predicting RNA tertiary structure formation and energetics.

## INTRODUCTION

Structured RNAs perform a wealth of essential biological functions, including the catalysis of peptide bond formation, gene expression regulation, and genome maintenance. In each case, the RNA folds into a complex three-dimensional structure whose thermodynamics governs its function (1–5). Interrogation of the folding process has yielded general principles and has established that RNA structure forms hierarchically (6, 7). Extensive *in vitro* measurements of diverse RNA elements have enabled a thermodynamic model for the first step of RNA folding—RNA secondary structure formation—that can generally predict any RNA sequence (8, 9). On the other hand, no thermodynamic model exists to predict tertiary structure formation from secondary structure, despite the fact that this second step is fundamental to RNA function.

RNA tertiary structures are composed of helices, junctions, and more sparsely distributed tertiary contacts (10). During tertiary folding, tertiary contacts form between distal interfaces, but only when brought into close enough proximity (11–13). Predicting this likelihood relies on understanding the conformational preferences of each of the RNA elements separating the interaction interfaces. To date, the primary focus in RNA tertiary structure folding has been on non-canonical motifs as their topology makes their structure challenging to predict and are thought to impart most of the conformational flexibility (14–22). However, within structured RNAs, over 50% of residues are contained in Watson-Crick (WC) base paired helices (23), implying that even subtle conformational flexibility in WC base pairs (as observed in (24–26)) might accumulate to substantially influence tertiary structure folding. Indeed, the 3D structure of RNA helices is known to depend on their sequence composition. For example, distinct RNA helices have different mechanical properties (24) and distinct chemical shift profiles, as determined in Nuclear Magnetic Resonance (NMR) (27). Whether these sequence-dependent structural preferences would affect the process of tertiary structure formation has not yet been tested.

Here, we have dissected the effects of helical sequence and flexibility on tertiary structure formation by combining two experimental and computational approaches that are each massively parallel and highly quantitative. Experimentally, we harnessed a recently developed platform to measure the association of structured RNAs at high-throughput, allowing both massively parallel and quantitative measurements of the formation of numerous model RNA heterodimers (tectoRNAs; Figures 1A-B) (28–31). Computationally, we collated the sequence-dependent conformations of WC base pair steps based on the existing X-ray crystal structure database and used this conformational variation to build a thermodynamic model for tertiary assembly. Computational simulations generated blind predictions of the relative affinity of all possible sequence-changing and length-changing helical variants of one piece of the tectoRNA heterodimer (>10^5^ predictions). We then measured >1500 of these previously uncharacterized tectoRNA variants. Our results establish that sequence-dependent conformational effects of helical elements can influence the thermodynamic stability of tertiary structures over an unexpectedly wide range of several kilocalories per mole, and that these effects can be predicted through computer modeling with surprisingly quantitative accuracy.

**Figure 1.**
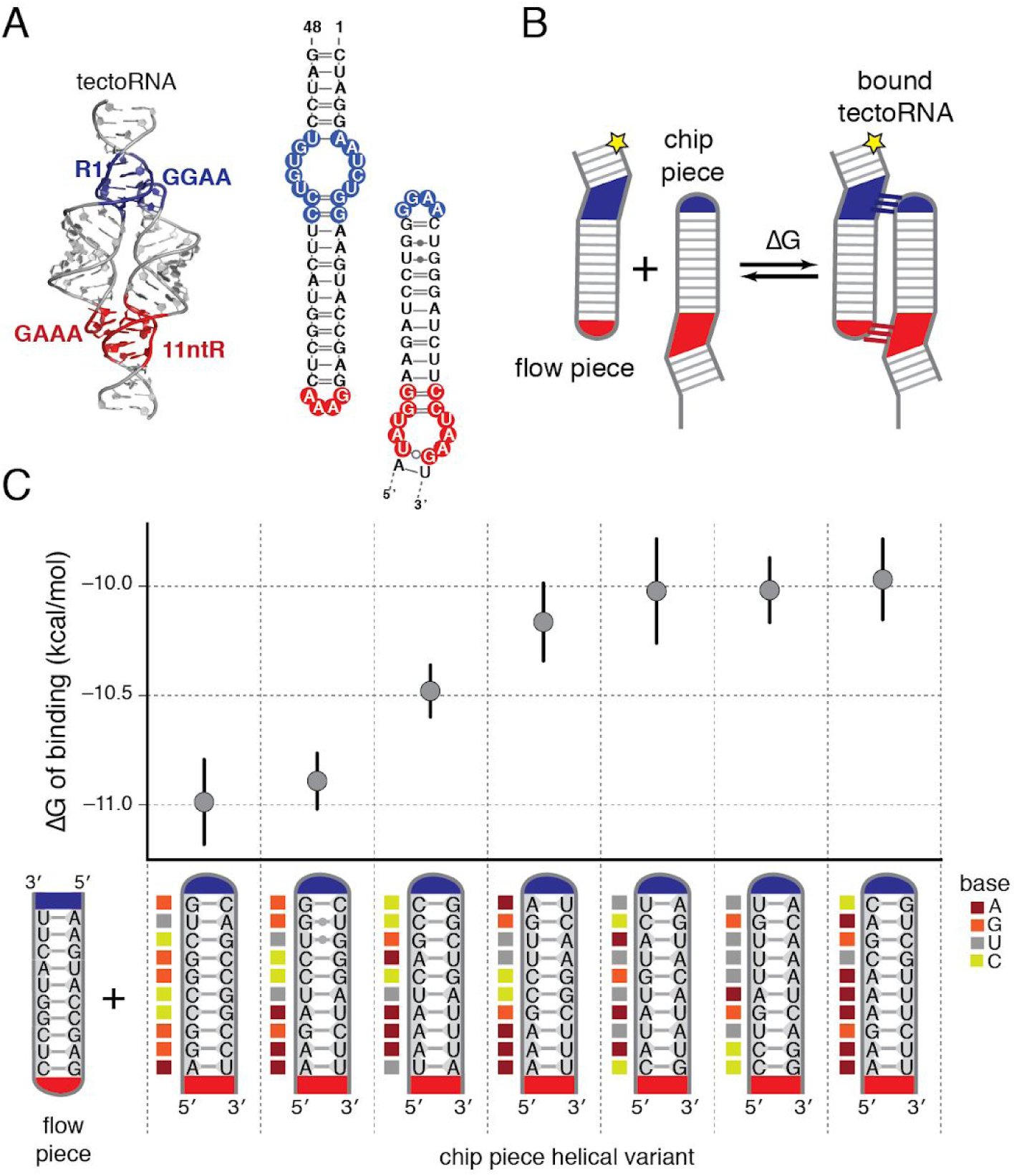
Free energy of tectoRNA binding depends on helix sequence. A) Structure of tectoRNA homodimer (PDB: 2ADT) with two tertiary contacts (GAAA-11nt). One of these tertiary contacts is replaced (GGAA-R1; blue) to convert the complex to the heterodimer used in this study (49). At right is the sequence and secondary structure of the wild-type tectoRNA interaction. Numbers indicate the “position” within the chip piece helix. (B) In our experimental setup, one piece of the heterodimer was fluorescently-labelled and free in solution (the “flow piece”), while the other was immobilized on the surface of a sequencing chip (“chip piece”). Quantification of bound “flow” piece to the chip surface allowed determination of the free energy of binding (ΔG) to form the bound tectoRNA. (C) Free energy of binding of the flow piece to seven distinct chip piece variants. Error bars are 95% CI on the measured ΔG. The sequence of the flow and chip piece helices is indicated (bottom).

## RESULTS

### High-throughput platform to measure thermodynamic stability of tectoRNA

To probe the effect of RNA helix composition on the formation of RNA tertiary structure, we used tectoRNAs as a test system (32). The tectoRNA is a heterodimer composed of two structured RNAs and contains two intermolecular tetraloop/tetraloop-receptor (TL/TLR) tertiary contacts (Figure 1A). The likelihood of forming both contacts depends on the conformational preferences of the RNA helices that bridge the two tertiary contact interfaces. Thus, measuring the thermodynamic stability of forming the tectoRNA assembly provides a quantitative readout of the conformational preferences of its constituent base pairs. Furthermore, the simple architecture of this system (with only ten base pairs in each of the two helices) enables investigation of the sequence effects on tertiary structure formation.

To profile affinity of tectoRNA heterodimers at high throughout, we designed a library of variants of one of these structured RNAs, the “chip piece” (Figure 1B). These variants were synthesized as DNA templates and amplified to include sequencing adapters and regions for RNAP initiation (Supplemental Figure 1A). This library of variants was sequenced; after sequencing, each DNA variant was *in situ* transcribed into RNA (Methods), enabling display of sequence-identified clusters of RNA on the surface of the sequencing chip (Supplemental Figure 1B) (30). The fluorescently-labeled tectoRNA binding partner, the “flow piece”, was introduced to the sequencing chip flow cell at increasing concentrations, allowing quantification of bound fluorescence to each cluster of RNA after equilibration (Methods). These fluorescence values were used to derive the affinity of the flow piece to each chip piece variant, in terms of the dissociation constant (*K*_d_) and binding free energy, (ΔG = *RT* log(*K*_d_)). Values for ΔG obtained in two independent experiments were highly reproducible (R^2^ = 0.92; RMSD = 0.15 kcal/mol; Supplemental Figure 2A). Each chip piece variant was present in multiple locations per chip (N ≥ 5), allowing estimation of confidence intervals for each affinity measurement (median uncertainty on ΔG = 0.16 kcal/mol (95% CI); Supplemental Figure 2B).

A preliminary experiment measured seven chip-piece RNA variants with different WC base pair compositions. We observed a range of binding affinities of 5 fold (1 kcal/mol; Figure 1C), demonstrating a relationship between the stability of tectoRNA assembly formation and the sequence composition of its helices. This observation inspired the development of a predictive computational model (described below) to relate helix structure to tectoRNA stability, based on structural differences between WC base pairs.

### Computational model for tectoRNA stability using conformational ensembles of RNA helices

Our computational model for tectoRNA stability relied on modeling the conformational ensemble for each RNA helix sequence—i.e., the distribution of conformations that the unconstrained helix explores in solution. Inspired by previous modeling procedures pioneered by Olson and colleagues (see (24, 33)), we divided each helix into the set of base pair steps (i.e., two sequential base pairs) (Figure 2A). Decomposition of helices in this manner allows for modeling of arbitrary helix sequences using a minimal set of structural states. Base pair step conformational ensembles were determined by collating the RNA crystal structure database for all instances of that base pair step in any structured RNA (Figure 2A, right) (24, 34), (35, 36). These structures were then clustered based on structural similarity to form a set of 50–250 discrete conformational states, each weighted according to its frequency (Methods and SI Table 1). The model takes advantage of RNA structural and ensemble modularity, the hypothesis that each RNA motif has its own set of conformations that together convolve into the global state of an RNA (12, 31, 37, 38).

**Figure 2.**
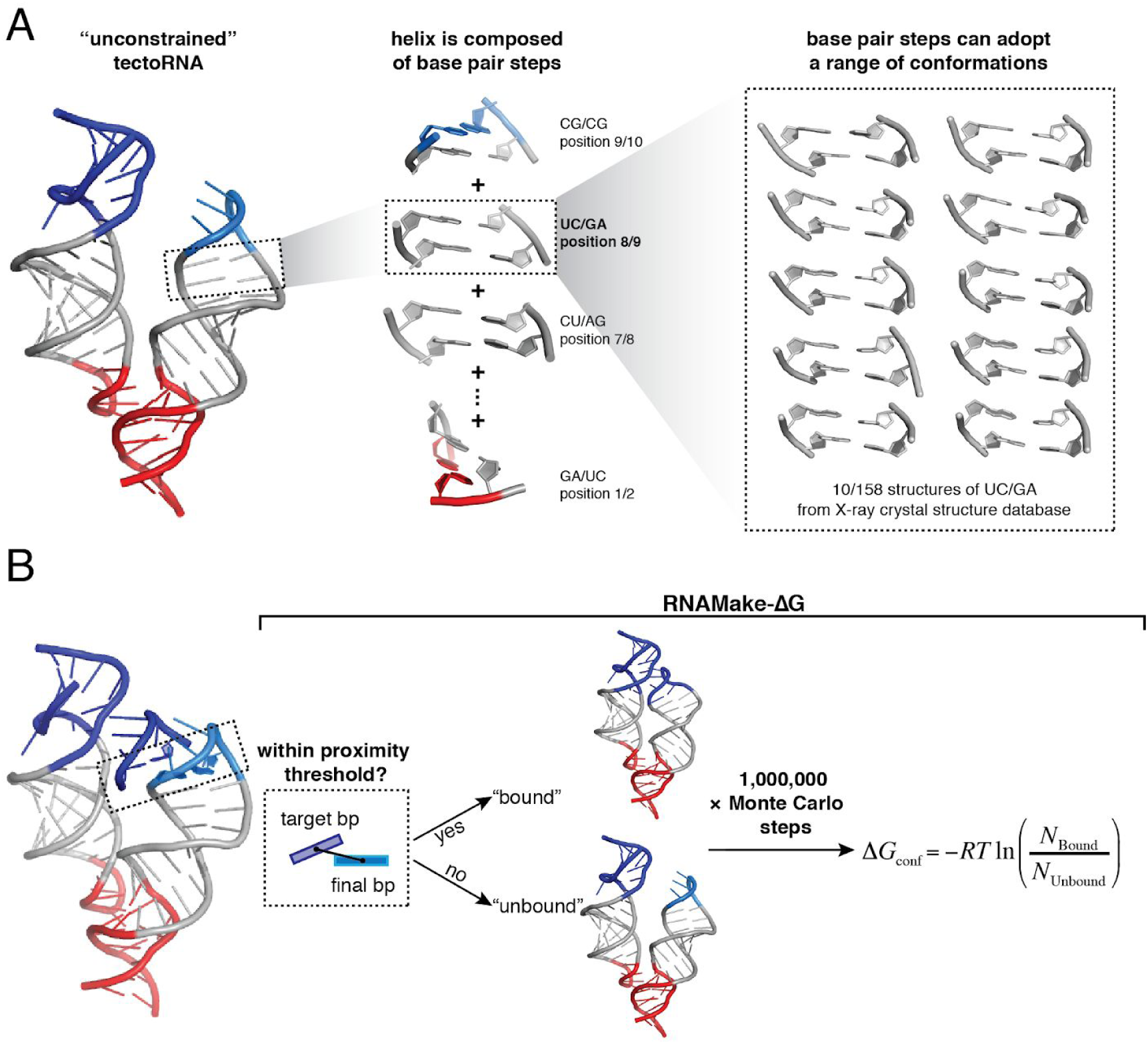
Ensemble model for RNA helices allows prediction of tectoRNA assembly energetics. A) (Left) The modeled structure of the unconstrained tectoRNA (i.e. with one contact formed) is shown. The global structure was assembled from the structures of its constituent elements, including the base pair steps that compose the helical regions. (Center) Example base pair steps are shown for the chip piece helix. Each base pair step can adopt an ensemble of many possible conformations, which were derived from examples of that base pair step in the crystallographic database. (Right) Example conformations within the UC/GA conformational ensemble are shown. B) Starting with the unconstrained tectoRNA as shown in (A), a Monte Carlo simulation was performed. At each step of the simulation, the structure of one base pair step in the tectoRNA was replaced with a new state from its conformational ensemble. The new structure of the unconstrained tectoRNA assembly was evaluated for whether it was “bound” or “unbound,” according to the translational and rotational distance from the target base pair to the final base pair. A million steps were performed and the total number of computed bound and unbound tectoRNA conformations were used to calculate the free energy change between the bound and unbound tectoRNAs (ΔG_conf_).

With this model we generated the “unconstrained” tectoRNA—i.e., the intermediate state of tectoRNA binding in which only a single tertiary contact is formed (Figure 2A). In this “unconstrained” state, the helices explore their full conformational ensembles (other than steric clashes) and occasionally bring the loop and receptor of the second tertiary contact in close enough proximity to form the closed tectoRNA assembly (Figure 2B). The relative population of conformational states that allow productive binding versus those that do not enables calculation of free energy penalty of forming the bound complex. This model was built as an extension of the RNAMake (a toolkit for the design of RNA 3D structure) (39) to predict thermodynamics of tertiary structure formation; thus we called the method “RNAMake-ΔG”.

Modeling the “unconstrained” tectoRNA additionally required structures for each of the TL/TLR tertiary contacts. Each TL/TLR receptor was modeled as a single structural conformation as opposed to a conformational ensemble, as this type of tertiary contact appears nearly structurally identical across all extant crystallographic structures (12). The single structure of the GAAA-11nt receptor interaction was derived from the crystal structure of the P4-P6 domain of the Tetrahymena group I intron (PDB 1GID), and the structure of the GGAA-R1 interaction (R1) was derived from the Rosetta stepwise Monte Carlo method (40).

A Monte Carlo simulation was employed to assess the conformational behavior of the unconstrained tectoRNA. During each step of the simulation, a new conformation for one of the base pair steps within the assembly was sampled from the base pair step’s conformational ensemble—this altered base pair structure was used to recalculate the overall structure of the unconstrained tectoRNA. The simulated tectoRNA was assessed for whether it could form the second contact, closing the assembly: The position of the closing base pair of the unbound tetraloop in the unconstrained tectoRNA was compared to its position in the bound TL/TLR (Figure 2B). A proximity threshold of 5 Å and a rotational alignment term (see Methods and refinement below) was imposed on this position difference to define whether the structure was closed with both contacts formed (“bound”) or not (“unbound”) (Figure 2B). These values were used to calculate the free energy of conformational alignment of the tertiary contacts:

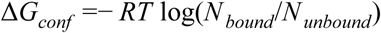

where T is the temperature, R is the universal gas constant, and *N_bound_* and *N_unbound_* are the number of simulated structures called as bound or unbound, respectively. We attributed differences in binding affinity between any two tectoRNA variants (ΔΔG_binding_) to differences in this conformational alignment term:

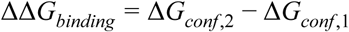

where ΔG_conf,1_ and ΔG_conf,2_ are the conformational alignment terms for two variants (indicated by 1 and 2, respectively). We generated ΔG_conf_ for all possible sequences of the four canonical base pairs within the chip piece helix (Methods). These RNAmake-ΔG calculations suggested that changes in helix sequence could change the predicted ΔG of tertiary binding over a substantial range of 2.5 kcal/mol, corresponding to a 70-fold effect on affinity (Supplemental Figure 3). We test this model below.

### Blind prediction of RNA assembly energetics with RNAMake-ΔG

We next tested the predictions of RNAMake-ΔG in a blind prediction challenge. We selected 2000 tectoRNA sequences that were predicted (by author JDY) to uniformly span the predicted range of affinity. Two authors (SD and NB) then carried out high-precision measurements for 1596 of these sequences (the remaining 404 sequences were not sufficiently represented in our library). The tested sequences gave experimental tertiary stabilities spanning a range of affinity of 2.1 kcal/mol (corresponding to a 40-fold effect on *K*_d_) between the lowest and the highest affinity binders, similar to the 2.5 kcal/mol (70-fold effect) predicted range. These data confirmed that sequence-dependent conformations of RNA helices can have a substantial effect on tertiary structure formation. Strikingly, we observed a high correlation between the observed and predicted affinities (R^2^ = 0.71), with RMSD of 0.34 kcal/mol to the predicted line of fixed slope = 1 (Figure 3A). Allowing the slope to vary gave a slightly better prediction (RMSD = 0.21 kcal/mol; best-fit slope = 0.54) (Figure 3A). The good agreement between our observed and predicted values suggests that this empirical model captures important structural differences among helices that in turn determine the thermodynamics of tertiary structure formation.

**Figure 3.**
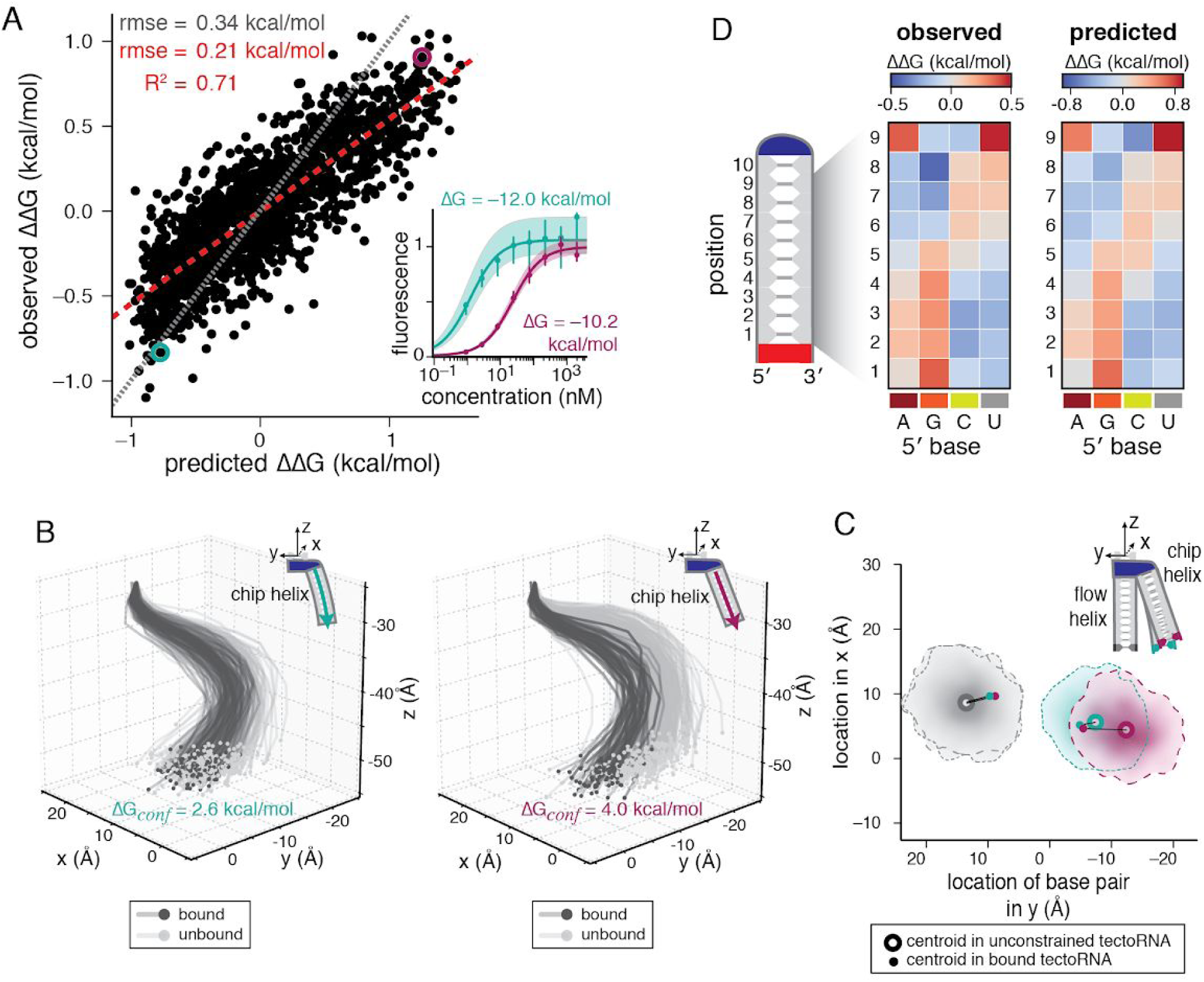
RNAMake-ΔΔG accounts for changes in tectoRNA affinity in a blind prediction challenge. A) Scatterplot compares the dependence of the observed values of tectoRNA binding ΔG on the predicted values generated with the RNAMake-ΔG model, for 1536 chip piece variants (R^2^ = 0.71). Each set of ΔG values is compared to their respective medians to obtain ΔΔGs. Red dashed line indicates the best-fit line (slope = 0.54); grey dotted line indicates the line of slope 1. Inset shows the measured binding affinity curves of two chip piece variants. B) Shown are example 3D trajectories of the chip-piece helix produced during the Monte Carlo sampling for two variants whose binding curves are shown in A). For each variant, 250 “unbound” trajectories (light grey) and 100 “bound” trajectories are shown (dark grey). All trajectories are aligned by the top base pair. C) Distribution of the terminating base pair of the chip and flow piece in the partially bound tectoRNA projected in the x-y plane. Distributions were determined using bivariate kde smoothing of ~1000 bound or partially bound structures sampled from the simulation. The centroids of the distributions are shown as open circles; black lines connect the centroid of the partially bound structures open circle to the centroid of the bound structures (black dot). D) Observed (left) and predicted (right) affinity for chip piece helices with the indicated base pair at each position within the helix. Affinities are given as deviation from the median observed or predicted affinity across all 1536 variants.

To help visualize the formation of the tectoRNA assembly, we compared the modeled conformational ensembles of two tectoRNA variants from the extremes of tectoRNA affinity (magenta = –10.2 kcal/mol, cyan = –12.0 kcal/mol; Figure 3A shows a subset of the chip piece helix trajectories, while Figure 3C shows the distribution of the final base pair of the flow and chip piece helix, projected in the *x*-*y* plane). Both the low- and high-affinity chip piece helices sampled a wide range of RNA backbone trajectories in the unconstrained tectoRNA, with variation in the position of the final base pair of more than 7 Å (full width at half maximum in *x*- and *y*-directions; Figure 3C). The median position of the final base pair differed by 5.3 Å between the two chip piece helices, with the end of the helix being substantially farther from the flow piece for the low-affinity variant (Figure 3C). Tus, this variant was bound only in the subset of conformational states making more extreme conformational excursions (i.e., compare black and grey trajectories in Figure 3B), which results in the observed destabilization. These observations suggest that the helices of destabilized variants are bound in less probable, bent conformations that “reach” to form the tertiary interaction.

Attempting to model binding affinity using only a single, most likely structure for each base pair step produced worse predictions (R^2^ = 0.42), highlighting that the flexibility of base pair steps cannot be ignored in assessing thermodynamics of tertiary structure formation (Supplemental Figure 4A-B). We observed that certain structural differences had large effects on thermodynamic stability, while others had minimal effects (Supplemental Figure 4C-D & next section), an observation we could not have made without our ensemble model. By taking the difference between the centroid of the bound states and unconstrained states, we determined a projection of the structural differences most coupled to thermodynamic effects (Methods). Differences between helices along this project were highly correlated to the observed ΔΔG values (R^2^ = 0.71; Supplemental Figure 4D), while differences along a perpendicular axis were uncorrelated. Thus, certain differences between single structures may be used to predict thermodynamic effects, but this prediction required making additional assumptions for how structure differences relate to thermodynamic differences, in contrast to a full ensemble model that allows direct calculation of thermodynamic parameters.

### Base pair elements adopt distinct structures at different positions

To gain insight into the how primary sequence affects binding probability in this system, we determined the average effect on tectoRNA affinity (ΔΔG) of having any given base pair at each position within the helix, compared to the average affinity of all 1594 tested variants (Figure 3D; Supplemental Figure 5B-C). These effects were highly correlated between the observed and predicted values (*R^2^* = 0.93; Supplemental Figure 5A), and this agreement is particularly striking in a heat-map representation (Figure 3D). Interestingly, each base pair can have either stabilizing or destabilizing effects depending on its position within the helix (Figure 3D). Base pairs with a purine residue on the 5’ side of the helix (i.e., A-U and G-C base pairs) were destabilizing when placed closer to the receptor (positions 1–3), but stabilizing when placed closer to the loop (positions 6–8), while the reverse was true for base pairs with a purine on the 3’ side of the helix (i.e., U-A and C-G base pairs; Figure 3D). This observed position-dependence of sequence preference strongly contrasts with the ‘nearest-neighbor rules’ governing secondary structure energetics, in which each base pair step contributes an additive free energy term toward the overall free energy of folding, regardless of its position within a helix (41), suggesting that partial unfolding of the secondary structure is not responsible for the differences in tectoRNA assembly formation (see Supplemental Figure 6).

The overall trend in position dependence suggests a simplifying rule that conformational preferences of purine-pyrimidine base pairs are similar, but are distinct from pyrimidine-purine base pairs. However, an exception to this rule is evident at position 9, where A-U and U-A were both destabilizing. This base pair is adjacent to the closing base pair of the loop, leading us to consider whether this base pair adopted substantially different conformations in the bound tectoRNA due to the proximity of the tertiary contact. However, the observed effect was highly correlated with the effect predicted by the RNAMake-ΔG model (Figure 3D, “position 9” row), suggesting that the loop-proximal base pair step is not appreciably sampling conformations absent from the conformational ensemble used in RNAMake-ΔG.

The position-dependent effects might arise from a preference for different structural orientations at each position within the helix of the bound tectoRNA. We characterized the difference in the average structural coordinates of each base pair step element in the bound tectoRNA ensembles and the unconstrained tectoRNA ensembles (Methods). The resulting structural differences were slight (<0.3 Å in any of the translational coordinates) but position-dependent, supporting this hypothesis (Supplemental Figure 7). The magnitude of position-dependent structural differences is consistent with each position exerting a small contribution toward achieving an overall bent conformation. Understanding the overall energetic effect therefore requires understanding each of these small base pair step contributions.

To achieve a more granular understanding of the position-dependent structural preferences of base pair steps, we quantified the contribution of each of the base pair step’s conformational states in the bound tectoRNA. States with an increased representation (over and above the expected sampling frequency from the Monte Carlo simulation) in the bound tectoRNA should correspond to the states that promote binding, and vice versa for those with a decrease in representation (Supplemental Figure 7). We observed disproportionate representation of certain substates within each base pair step’s ensemble in the bound tectoRNA (illustrated for the AU/AU ensemble in Figure 4A and for all base pairs in Supplemental Figure 8). Notably, these changes were highly position dependent, such that the majority of states could be overrepresented or underrepresented, depending on their position within the chip piece helix (Figure 4A). For the AU/AU ensemble, conformational states were clustered based on their position-dependent representation (shown in dendrogram and colors in Figure 4A). Conformational states in different clusters were each associated with distinct structural behavior, with >1 Å translational differences between structures in different clusters (Figure 4B-C). For example, conformers in class 6, which promote binding in positions 1–3 in the helix, are more twisted and thus span less translational distance than conformers in class 1, which promote binding only in the very first or last base pair in the helix (Figure 4B-C). These results suggest that the same base pair element adopts substantially different conformations depending on its location within the helix, thereby accounting for the differential base pair preferences along the helix (Figure 3D). These different conformational preferences further underscore the necessity of an ensemble to account for thermodynamic effects in this system.

**Figure 4.**
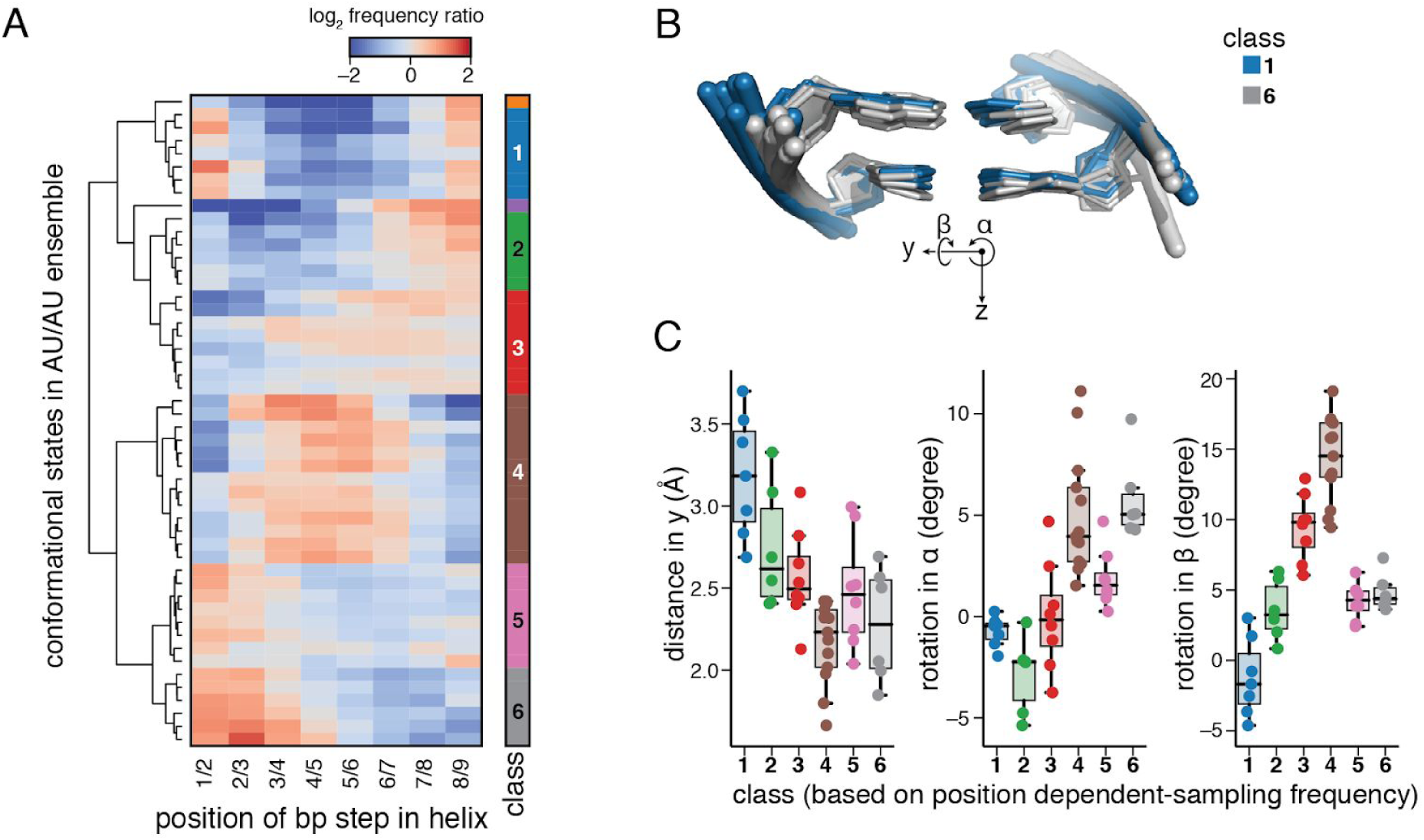
Ensemble model is required for accurate energetic predictions. A) Change in sampling frequency of conformational states in the AU/AU ensemble in the bound versus the partially bound. B) example structures of base pair step conformations that are enriched and depleted at two positions. C) Change in positioning between enriched and depleted conformational states at each position of any base pair type. (see Supplemental Figure 8 for other coordinates). Enriched = sampling frequency more than 2 fold greater than expected, depleted = sampling frequency less than 2 fold less than expected.

### Testing RNAMake-ΔG at more extreme helical distortions

While RNAMake-ΔG was surprisingly accurate at predicting the effects of sequence-dependent helical conformations on tectoRNA assembly formation of two 10 bp helices, we next aimed to explore the limits of the predictive power of RNAMake-ΔG by adding or deleting base pairs on both the flow and chip RNAs. We generated chip RNAs with helix lengths of 8 bp to 12 bp (N = 32 to 96 sequence variants per length) and tested each chip RNA against flow RNAs with helix lengths of 9 bp to 11 bp, yielding 15 length pair combinations (Figure 5A). Each of these complexes was destabilized relative to the original assemblies with 10 bp flow and 10 bp chip helices, which we abbreviate “10/10 bp”. Certain highly mismatched length combinations were so destabilizing that no binding was detectable (ΔΔG > 4.4 kcal/mol relative to 10/10 bp; 8 length pair combinations; Supplemental Figure 9). The remaining length pair combinations had effects spanning a 4.4 kcal/mol range. The thermodynamic stability of each length pair complex with observable binding was calculated with RNAMake-ΔG. Comparisons to measurements demonstrated a correlation of R^2^ = 0.66 and RMSD = 0.72 kcal/mol for these predictions, with the best-fit line having slope indistinguishable from one (Figure 5A). The larger RMSD compared to the 10/10 bp sequence predictions appears due to certain length pairs exhibiting systematic deviations between the observed and predicted effect. For example, the 10/11 bp flow/chip complexes were uniformly observed to bind more weakly than predicted, while the 9/9 bp flow/chip complexes were observed to bind slightly tighter than predicted.

**Figure 5.**
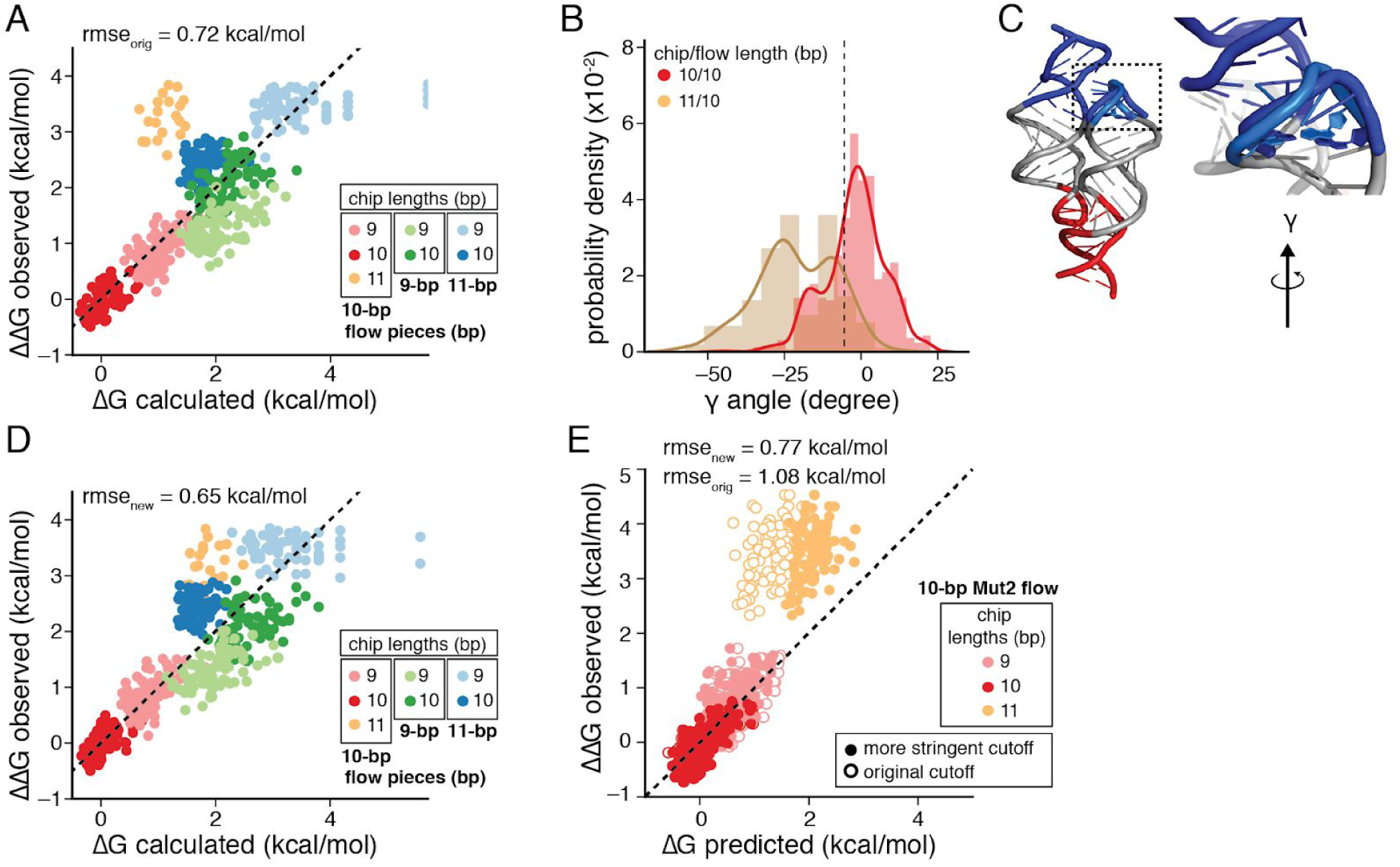
Increased prediction accuracy of different length pairs with refinement of the bound state cutoff. A) Scatterplot of the observed vs calculated affinity for chip- and flow-piece variants with altered lengths. Colors indicate the length of the flow- and chip-piece helix. B) Distribution of the value of for gamma within two bound tectoRNA complexes, where gamma represents the rotation between the final bp and the target bp around the z-axis. The 11-bp chip piece variant has distinct values for gamma compared to the 10-bp chip piece variant. Vertical dashed line indicates γ = –10 degrees. C) Structure of the bound complex with the original cutoff (light blue) or a more stringent cutoff (blue) where γ has to be > –10 degrees. D) Scatterplot of the observed and calculated affinity for length pair combinations with the more stringent cutoff which excluded overtwisted conformations (i.e. γ > –10 degrees in the bound complex) Colors indicate the length of the flow- and chip-piece helix, as in (A). Observed values are the same as in (A). D) Scatterplot showing the observed vs predicted (blind prediction values) of a new set of chip piece sequences, against a distinct 10-bp flow piece, using either the original model (open circles) or the updated model with the more stringent cutoff (closed circles).

One possible explanation for observing a larger destabilization on binding than predicted is an overly accommodating proximity threshold for determining bound tectoRNA structures during prediction. Such a loose threshold would allow unrealistic structures to be considered bound during the RNAMake-ΔG simulation. To assess this possibility, we analyzed the predicted ensemble of bound tectoRNA conformations of a 10/11 bp flow/chip complex. In comparison to a bound 10/10 bp complex, there was a change in the twist of the base pair closing the tetraloop, leading to an underwound helix in the 10/11 bp flow/chip complex relative to the other length complexes (Figure 5B). We hypothesized that these states were binding incompetent, leading to the discrepancy between observed and predicted values for these length pair complexes (Supplemental Figure 10). To avoid classifying these states as bound, we tested a more stringent cutoff, implementing the additional criteria that the helix within the bound complex cannot be undertwisted (γ > –10°; see Figure 5E). With this additional constraint, the agreement between our calculated and observed ΔΔG for all length pairs improved significantly (R^2^ = 0.71; RMSD = 0.65 kcal/mol; Figure 5C).

To test this refined proximity threshold, we carried out a second blind prediction challenge, with calculations and experiments carried out independently by authors JDY and SD, respectively. The affinity of an additional 300 chip variants of three different lengths (9, 10, and 11 bp) were measured against a distinct 10-bp flow piece. Overall, these tectoRNA variants represented a wider diversity of sequences than those used to refine the proximity cutoff. The blind predictions using the additional constraint demonstrated a significantly improved relationship between the observed and predicted binding affinities, although it did not completely account for the destabilizing effect of this length pair complex (RMSE, original model = 1.08 kcal/mol; RMSE updated model = 0.77 kcal/mol; Figure 5D). The development of this additional constraint on the bound conformation demonstrates an iterative protocol for refining the anisotropic binding landscape of a tertiary contact.

## DISCUSSION

Structured RNAs form dynamic tertiary assemblies critical to cellular functions like translation and transcriptional regulation (1–3). A major goal in understanding these biological assemblies has been to develop a model for RNA structure formation from primary sequence. We have presented here a model for how RNA double helix flexibility can impact RNA tertiary structure energetics. The computational model is based on conformational ensembles of RNA double helices prepared by stitching together conformations of Watson-Crick base pair steps observed in the RNA crystallographic database. The model gives quantitative estimates for how sequence and length changes of helices change the favorability of bringing together segments that make RNA-RNA tertiary contacts. High-throughput measurements allow rigorous tests of this model and confirm its predictions with accuracies of 0.34 and 0.77 kcal/mol, for sequence and length changes, respectively. These accuracies are well under 1 kcal/mol, which is often considered the limit of computational modeling (so-called ‘chemical accuracy’). The agreement of computation and experiments is further demonstrated by comparisons of position-by-position sequence preferences. Finally, our modeling gives new physical description of how RNA helices ‘look’ inside tertiary assemblies. For example, the same base pair sequence is predicted to have slightly different physical structures when embedded at different position in the tertiary assembly, and this phenomenon explains the qualitatively different sequence preferences at each position. Furthermore, the model gives a view of such structural effects as spread throughout the helix and not focused at one particular “kink” within the helix. To rigorously make these conclusions and comparisons, it has been important to separate model building and model evaluation into different research groups, with the results confirmed through blind challenges. We envision that the rise of fast, high-throughput experimental measurements may enable broader adoption of the blind prediction paradigm, as implemented in this work.

Our work has strong implications for the biophysical modeling of RNA 3D structure. Historically, modeling RNA structure has been facilitated by the principle of RNA modularity—that RNA elements have transferable properties that can predict their behavior across diverse contexts. This principle underlies RNA secondary structure prediction, which relies on an independent and additive free energy contribution of each RNA base pair step to the overall secondary structure fold (8, 42). For RNA tertiary assembly, however, we observed that each base pair step could have opposite effects on the stability of the assembly when located in different positions within the helix, implying that a base pair step’s effect on tertiary structure stability is not a constant, transferable property. This indirect link between sequence information and thermodynamic effects may explain why the effect of helix sequence on tertiary structure formation has been difficult to discern prior to this study. Nevertheless, mitigating this complexity, the success of our model at predicting these context-dependent thermodynamic effects ultimately supports the principle of RNA modularity, but it is the base pair step’s conformational ensemble that is transferable, not its energetic contribution. We anticipate that the insight of *ensemble modularity* applied within our computational framework can predict the effect of helix sequence in diverse contexts, ultimately making it easier to discern contexts in which helix sequence is important in maintaining a specific tertiary conformation.

We envision that an improved RNAMake-ΔG can be extended to predict thermodynamic stability of more complex systems and to help determine the conformational ensembles of non-canonical motifs. Indeed, we have already experimentally examined the effects of non-canonical motifs, such as bulges, on the stability of the tectoRNA system (31). Refining the relevant conformational ensembles using increased crystallographic statistics, ab initio computer modeling, and high-throughput data is an important next challenge. Finally, we speculate that our algorithm will help in understanding the energetic costs and sequence preferences associated with RNA double helix contortions that occur throughout important biological processes ranging from tRNA-based decoding by the ribosome to the packaging of RNA inside viruses.

## METHODS

All software and source code used in this work are freely available for non-commercial use. RNAMake software and documentation are at https://github.com/jyesselm/RNAMake.

### Flow piece labeling

Three distinct flow pieces were used to probe the chip piece library, with helices of length 9, 10 (wildtype), and 11 base pairs (see Table 1). Flow pieces were ordered as RNA oligos from Integrated DNA Technologies (Coralville, Iowa) with a 5′-Amino Modifier C6 modification, with HPLC purification. Each flow piece was ethanol precipitated at –20 °C overnight, followed by resuspension to a final concentration of 2 mM with 2 mM of NHS-conjugated Cy3b dye in 50 mM phosphate buffer (pH 8.7). This reaction was incubated at 37 °C for 1 hour, followed by PAGE purification (8% PAGE, 8 M Urea, 1x TBE: 89 mM Tris-HCl, 89 mM Boric Acid, pH 7.4, 2 mM sodium EDTA). RNA was eluted from the gel in water using three freeze-thaw cycles. To reduce aggregation on the chip surface, flow piece solutions were spun in a 50K Amicon filter two times and collected on a 3K Amicon filter. Flow pieces were quantified after purification using Qubit RNA high sensitivity kit (Thermofisher).

**Table 1:**
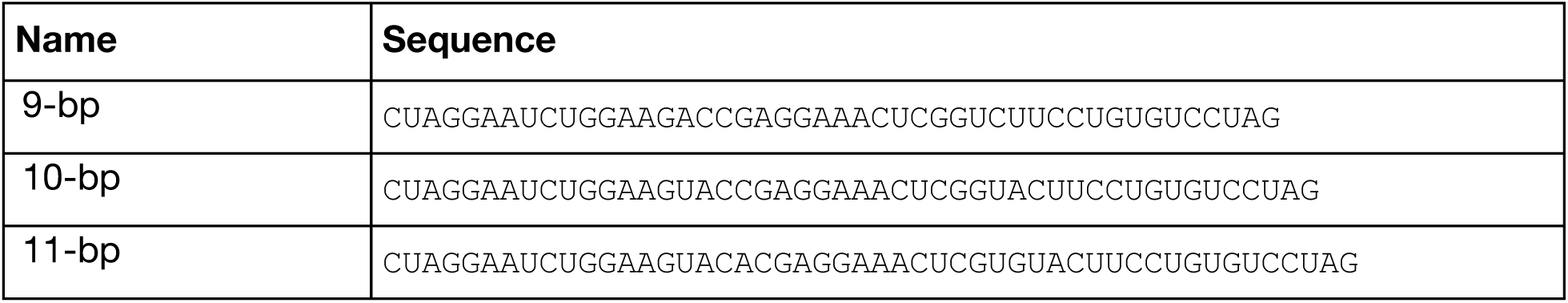
Flow piece sequences.

### Chip piece library design, amplification, and sequencing

The tectoRNA library was designed by replacing the chip piece helix with a set of defined WC base pair sequences. This library of chip piece variants (~2000 sequences) was ordered together with other tectoRNA variants not discussed here, to form a final library of ~45,000 variants. The library was ordered with common priming sequences across chip piece variants from CustomArray (Bothell, WA). This pool of DNA oligonucleotides was PCR amplified with primers oligopool_left and oligopool_right (see Table 2 and Supplemental Figure 1A), with 1:400 dilution of the synthesized oligo pool, 200 nM of each primer, 200 μM dNTPs, 3% DMSO, 1x Phusion HF buffer, 0.01U/μl of HS Phusion (NEB). Primers were purchased from Integrated DNA Technologies (Coralville, Iowa). The reaction proceeded for 9 cycles of 98 °C for 10 seconds, 62 °C for 30 seconds, and 72 °C for 30 seconds, followed by cleanup of the reaction mixture using Qiagen PCR Cleanup Kit (elution into 20 μl). To append sequencing adapters to this PCR product as well as include unique molecular identifier (UMI, in the form of a 16 nt random N-mer), a five-piece PCR assembly was performed, with 1 μl of the previous reaction, 137 nM of primers (short_C and short_D; Table 2), 3.84 nM of the adapter sequences (C1_R1_BC_RNAP and D_Read2; Table 2), 200 μM dNTPs, 3% DMSO, 1x Phusion HF buffer, and 0.02U/μl of Phusion Hot Start Flex enzyme (NEB). The reaction proceeded for 14 cycles of 98 °C for 10 seconds, 63 °C for 30 seconds, and 72 °C for 30 seconds, followed by cleanup with Qiagen PCR Cleanup Kit, as above.

**Table 2:**
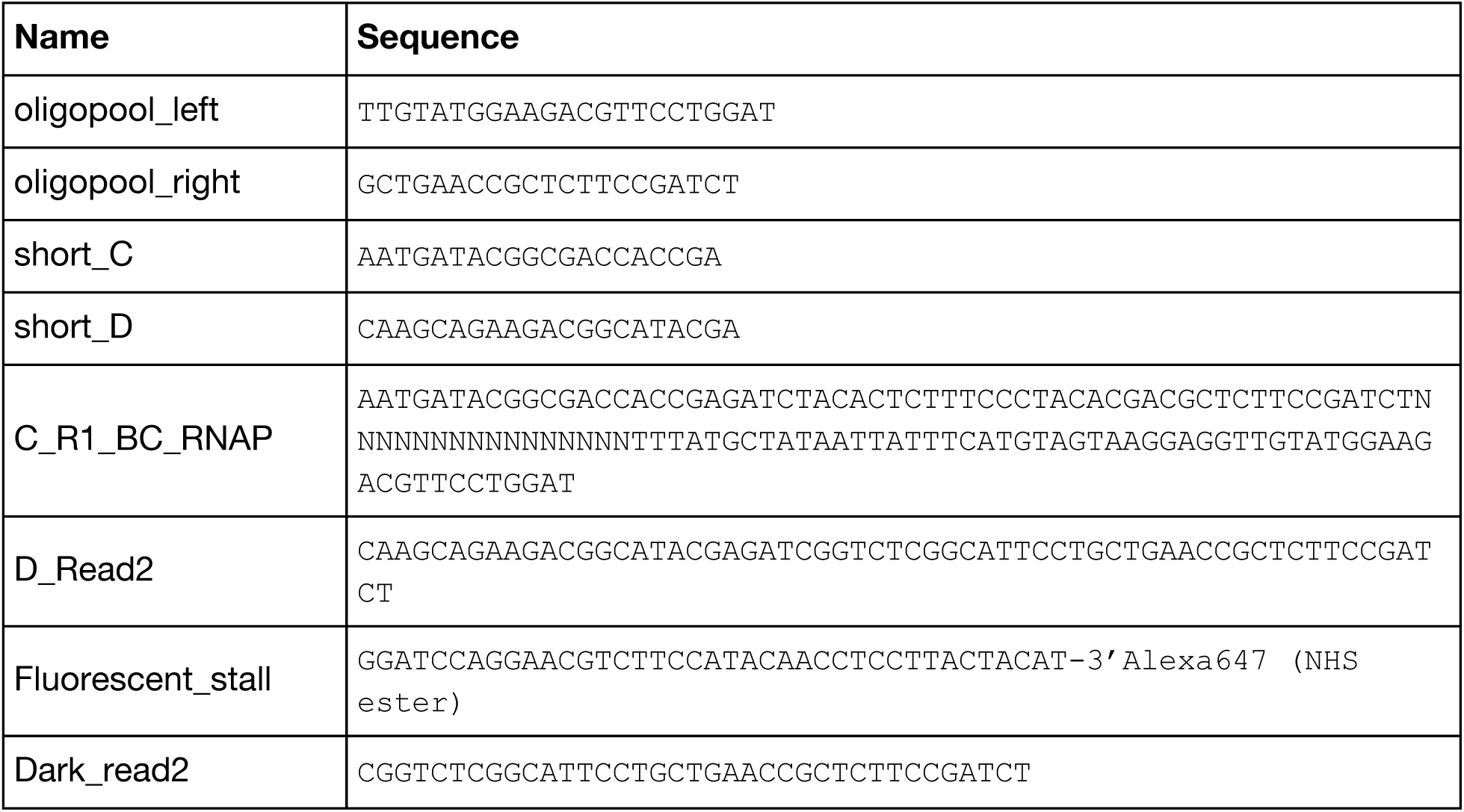
Primers used to amplify library for sequencing.

After amplification and assembly, the library was bottlenecked to reduce the representation of UMIs to ~700K distinct 16 nt N-mers. First, the library was diluted 1:5000 in 0.1% Tween20, and this dilution was quantified against a standard library of PhiX (Illumina, Hayward, CA), which was diluted two-fold seven times to form a dilution series from 25 pM to 0.2 pM. The standard series and the library dilution were amplified in a qPCR assay to determine their relative cycle threshold (CT) values; these values were used to determine the concentration of the diluted library by linear regression analysis of the CT values against the known concentrations of the standards. The volume associated with 700K molecules was PCR amplified, with 1.25 μM of primers (short_C and short_D), and 1x NEBNext Master Mix (NEB, M0541S), for 21 cycles of 98 °C for 10 seconds, 63 °C for 30 seconds, and 72 °C for 60 seconds, followed by cleanup with Qiagen PCR Cleanup Kit. The final library was sequenced on an Illumina Miseq instrument at 10-30% of the total sequencing chip, with the rest of the chip consisting of high-complexity genomic sequences. Sequencing cycles were performed as follows: 75 bases in read 1, 75 bases in read 2, and an 8 bp i7 index read, resulting in demultiplexed, paired-end sequences.

The output of the Illumina sequencing included the read1 and read2 sequence associated with each cluster ID. This information was processed to extract the UMI sequence from read1 for each cluster (by extracting the sequence preceding the RNAP initiation site; see Supplemental Figure 1A). Clusters with common UMI sequences were processed to obtain a consensus read2 sequence, by taking the most common base at each position (i.e. per-base voting consensus). UMIs of poor quality, with poor representation or poor agreement across sequences, were removed, by assessing the number of clusters with read2 sequences matching the consensus sequence. Poor quality was defined if the number of matches (or successes) could be explained by a null model with p value > 0.01, where the null model was a binomial distribution with probability of success of 0.25. This filter removed UMIs associated with diverse unrelated sequences, or with relatively few reads per UMI.

Finally, the consensus sequence of each UMI was associated with each designed library variant by searching for an exact match of the reverse complement of the designed sequence within the read2 consensus (starting at the first base).

### Experimental platform for parallel measurements on a sequencing chip

The sequencing chip used for Illumina Miseq sequencing was directly used on a custom-built imaging station, made from a combination of parts from an Illumina Genome Analyzer IIx and parts that were custom-designed, as described originally in (30), and modified as in (29). The flow cell surface was imaged with a TIRF setup, allowing measurement of the bound fluorescence on the chip surface with minimal background from fluorescent molecules in solution. Custom scripts were used to control the laser power, stage, temperature, fluidics, and camera. Images could be taken in one of two channels, the “red” channel (660 nm laser, with 664 nm long pass filter from Semrock), and the “green” channel (530 nm laser and a 590 (104) nm band pass filter from Semrock). To image the flow cell surface, 16 images were taken to overlap tiles 1 through 16 taken of the Miseq sequencing output. Each image was taken for 400 ms exposure time with 200 mW input laser power.

RNA was generated in situ on the surface of the Illumina Miseq chip by a series of enzymatic reactions carried out through fluidic application and temperature control, as described in (29, 30, 43). In brief, covalently attached ssDNA was converted to dsDNA through extension of a biotinylated primer, followed by incubation with streptavidin to create a streptavidin roadblock (see Supplemental Figure 1B). E. coli RNA Polymerase (NEB M0551S) was applied to the flow cell with limiting concentrations of NTPs (2.5 μM each of ATP, GTP, and UTP), allowing only very limited extension and preventing initiation by more than one polymerase per molecule. Excess polymerase was washed out of the flow cell, followed by incubation with the full suite of NTPs at high concentration (1 mM each NTP) to allow extension. Encountering the streptavidin roadblock causes polymerases to stall, resulting in stable display of the nascent transcript (Supplemental Figure 1B). Detailed descriptions of each of these steps may be found in (43).

After RNA extension, blocking oligos were annealed to common regions on the nascent transcript (see Supplemental Figure 1A) to limit the formation of alternate secondary structure, as well as to fluorescently label clusters of transcribed RNA (fluorescent_stall and dark_read2; Table 2). Oligos were purchased from Integrated DNA Technologies (Coralville, Iowa) with RNase-Free HPLC Purification.

### On chip experiments to determine tectoRNA affinity

For each experiment, a fluorescently-labeled tectoRNA flow piece was serially diluted three-fold to form a concentration series from 2000 nM to 0.91 nM in binding buffer (89 mM Tris-Borate, pH 8.0, 30 mM MgCl_2_, 0.01 mg/ml yeast tRNAs (ThermoFisher Scientific AM7119), 0.01% Tween20. To fold the flow piece, it was initially diluted to 10 uM in water, and denatured by incubating for 1 minute at 95 °C, followed by refolding for 2 minutes on ice (preceding the dilution to 2 uM and serial dilution). Each flow piece solution was applied to the flow cell, and after waiting for sufficient time for equilibration, the flow cell was imaged in the red and green channels, with the red channel capturing the annealed oligo corresponding to any transcribed RNA, and the green channel capturing the bound flow piece. Experiments were carried out at at 22 °C. Equilibration times were as follows: 3 hours, 2 hours, 1 hour, 45 min, 30 min, 20 min, 20 min, and 20 min, for 0.91 nM, 2.7 nM, 8.2 nM, 25 nM, 74 nM, 222 nM, 667 nM, and 2000 nM, respectively. These times were calculated to allow equilibration for the most stable variants (i.e. ΔG of –12 kcal/mol or *K*_d_ of 1 nM), assuming a common association rate constant (*K_on_*) of ~6×10^4^ M^-1^ s^-1^(43).

### Quantification of ΔG from image series

Each image taken during the course of an experiment was processed to extract the fluorescence values of the Illumina Miseq clusters. First, the Miseq tile and x-y-positions of each sequenced cluster was determined (from the Miseq output). Because of differences in the optics of the Miseq and the imaging station, these coordinates did not correspond 1:1 to the pixel values of our images. To account for this, sequence data coordinates were scaled by an overall scale factor (of 10.96 imaging-station pixels to Miseq x-y position units). A global registration offset was determined by cross-correlation of the images and subsequent fitting of the cross-correlation matrix to a 2D Gaussian to obtain the x-y- position that maximized the cross correlation coefficient. Finally, to correct for nonlinear aberrations, this cross-correlation procedure was repeated for 256 subdivisions of the overall image to obtain corrections on the global x-y- position as a function of the location within the image. These corrections were fit to 2D surfaces for the x- and y- corrections, as a function of x- and y- position.

In each of the 256 subtiles, all clusters within the subtile were fit to a sum of 2D Gaussians, with x-y- positions given by the sequencing data coordinates, nonlinearly corrected as described above, as in (30). The integrated fluorescence associated with each cluster is then: 2π*A*σ^2^, where *A* is the amplitude and *σ* the standard deviation of the 2D Gaussian. The fluorescence associated with the bound flow piece was normalized by dividing by the fluorescence in the red channel, to account for variability of cluster size.

The series of concentration values for each cluster were fit to a binding isotherm, according tothe equation: 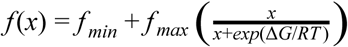, where *f* is the normalized fluorescence, *f_min_*, *f_max_*, and ΔG are free parameters, x is the concentration, *R* is the gas constant, and *T* is the temperature in Kelvin. Following single cluster fits, the values for *f_min_*, *f_max_*, and ΔG per variant were obtained by finding the median of these values across single clusters associated with each variant. An additional fitting step refined these values by applying a distribution for *f_max_* for those variants that did not achieve saturation, based on the values for *f_max_* of variants that did, as described in (43), ultimately allowing consistent attribution of the change in fluorescence values to changes in ΔG rather than *f_max_*. In brief, this fit refinement took the median fluorescence values across a set of clusters (resampled from all clusters associated with the variant). This set of median fluorescence values was fit to the binding isotherm equation, with *f_min_* set to the median fluorescence value across clusters that did not achieve saturation, and *f_max_* either allowed to float or set to a random value generated from the distribution of *f_max_*, depending on if the maximum fluorescence in the binding series did or did not exceed the lower bound of the 95% confidence interval of the *f_max_* distribution, respectively. This resampling and refitting was repeated 100 times for each variant, allowing determination of confidence intervals on the fit values of ΔG per variant.

### Combining experimental replicates

Data for the wildtype, 10-bp flow piece comes from two replicate experiments (shown in Supplemental Figure 2A). Values reported for this flow piece represent the average of the two replicate values, weighted by the inverse of the variance on each measurement. If the 95% confidence interval on ΔG is δΔ*G*, then the variance on the measurement is: σ^2^ =(δΔ*G*/1.96)^2^. Thus, the weighted average on ΔG is then: 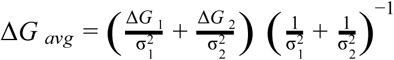. The combined error is then: 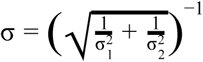.

### Building base pair step ensembles

To build a curated library of base-pair step components, we obtained the set of non-redundant RNA crystal structures managed by the Leontis and Zirbel groups (36) (version 1.45: http://rna.bgsu.edu/rna3dhub/nrlist/release/1.45). This set specifically removes redundant RNA structures that are identical to previously solved structures, such as ribosomes crystallized with different antibiotics. We processed each RNA structure to extract every motif using Dissecting the Spatial Structure of RNA (DSSR) (44) with the following command:

~~~
x3dna-dssr –i file.pdb -o file _dssr.out
~~~

We manually checked each extracted motif to confirm that it was the correct type, as DSSR sometimes classifies tertiary contacts as higher order junctions and vice versa. For each motif collected from DSSR, we ran the X3DNA find_pair and analyze programs to determine the reference frame for the first and last base pair of each motif to allow for alignment between motifs:

~~~
find_pair file.pdb 2> /dev/null stdout | analyze stdin >& /dev/null
~~~

We defined a base-pair step as two consecutive residues on one chain base paired to two consecutive residues on another chain, where both base pairs are in Watson-Crick orientation. Each instance of this pairing was collected from every structure. See Supplemental Table 1 for a summary of all total instances of each base-pair step.

### Clustering procedure for base pair step ensembles

To cluster the base-pair steps, all structures were first translated and rotated so that the first base pair was situated with its origin at (0,0,0) and its axes aligned with x, y and z orientation of the identity matrix, definition of base pair center and coordinate systems are as in (45). Fixed radius clustering was performed using a radius of a *distance*_*score* of 1.50, which was ideal according to optimization, although other radii did not greatly affect the final results. The *distance*_*score* between a cluster center and a new base-pair step is calculated below, where *d*_1_ and *R*_1_ are the translation and orientation of the cluster center’s second base pair, respectively. *d*_2_ and *R*_2_ are the translation and orientation of the second base pair in the base-pair step to be clustered.

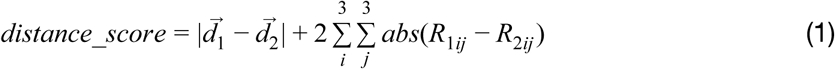

The number of clusters generated for each base-pair step sequence is shown in Supplemental Table 1. Each cluster was assigned a relative energy (Eq. 2) based on its population. *N_members_* is the number of base-pair steps in a given cluster, and *N_total_* is the number of base-pair steps of the current identity, i.e. AU/AU. This energy is used during our Monte Carlo simulations to allow swapping based on population.

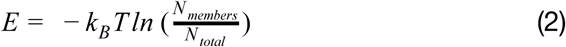

### TectoRNA simulation protocol

The simulation is set up by supplying a sequence and secondary structure for both tecto heterodimers. With this information, a 3D system is built up by representing each base-pair step with a corresponding structural ensemble and representing both tertiary contacts as single structures. The structure of the GAAA tetraloop/tetraloop-receptor (TTR) was isolated from the P4-P6 domain of the *Tetrahymena* ribozyme (PDB: 1GID). There is no known solved structure of the GGAA TTR; therefore, a structure was generated by modeling using stepwise monte carlo (40). The simulation proceeds by attempting to swap a randomly selected base-pair step from one conformation to another. If the new conformation has a lower energy, it is accepted; if not, it is selected by the metropolis criterion. All motifs are connected to each other by shared base pairs, so if a base-pair step is swapped from one conformation to another, the orientation change will propagate throughout the structure accordingly. In total, one million swaps are attempted during our standard simulation. To determine whether a conformation is bound, we calculate the distance_score (Eq. 1) between the final base pair of the chip helix and its original position (Figure 2A). If this score is lower than 5, we consider the conformation to be bound.

### Calculating the relative binding free energy of the tecto system

`rnamake_ddg` is part of a larger toolkit known as RNAMake. For instructions on installing RNAMake as well as extensive documentation available at http://jyesselm.github.io/RNAMake/. An example of running `simulate_tectos` is shown below.

~~~
rnamake_ddg \
-fseq “CTAGGAATCTGGAAGTACCGAGGAAACTCGGTACTTCCTGTGTCCTAG” \
-fsss “(((((( .... (((((((((((( .... )))))))))))) .... ))))))” \
-fcseq “CTAGGATATGGAAGATCCTCGGGAACGAGGATCTTCCTAAGTCCTAG” \
-css “((((((( .. ((((((((((((( .... ))))))))))))) ... )))))))” \
-s 1000000
~~~

The tecto system is composed of two distinct RNA molecules that dimerize. First is the “chip” piece, which is transcribed from the DNA on a MiSeq sequencing chip. There are up to one hundred thousand distinct sequences on each chip in a given experiment. The second sequence is the “flow” piece, which is titrated in during the experiment and can bind to all chip sequences. We maintain this nomenclature while running `rnamake_dg`. “`-fseq`” specifies the sequence of the flow RNA, and “`-fss`” specifies the corresponding secondary structure in dot-bracket notation. If a new sequence has the default secondary structure, “`-fss`” does not need to be used again. The flow sequence must include the GGAA tetraloop-receptor sequence and secondary structure or it will return an error. “`-cseq`” and “`-css`” are analogues to “`-fseq`” and “`-fss`”, but for the chip RNA. This RNA must include the GAAA tetraloop-receptor sequence or the output secondary structure will return an error. “`-s`” specifies the number of Monte Carlo steps to perform. The default is one million. The output of the program is the number of times that the Monte Carlo simulation sampled a “bound” conformation.

Using the output of the `rnamake_dg` program, it is possible to calculate the relative binding free energy of each sequence compared to the wild-type (WT) sequence where *N*_*bound* values are evaluated as the number of simulated conformations given distance score (eq. 1) compared to the target conformation of 5. Alternative forms of the distance score in (1), including more standard rotationally invariant metrics to define rotation matrix differences (46) or base-pair-to-base-pair RMSDs based on quaternions (47), but these were not tested in the current study.

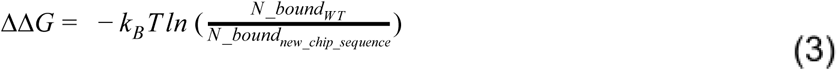

### Generation of 2000 helix sequences for blind predictions

To computationally assess the effect of the primary sequences of helices on relative binding, we generated all possible Watson-Crick helices. We put an A-U, U-A, G-C or C-G base pair at 9 positions in the chip sequence for a total of 4^9^ (262,144) sequences. For each generated sequence, we utilized RNAFold from ViennaFold (48) to confirm that the sequence folds into the target secondary structure. Then, we ran simulate_tectos on each new sequence with the following command.

~~~
rnamake_dg -cseq new sequence
~~~

## ACKNOWLEDGMENTS

We thank Curtis Layton and Johan Andreasson for developing, building, and maintaining the imaging station; Greenleaf and Herschlag lab members for reagents and critical feedback. This work was supported by the National Institute of Health (grant number P01 GM066275 to D.H., R01 GM111990 to W.J.G., R01 GM100953 and R35 GM122579 to R.D.). S.K.D. was supported in part by the Stanford Biophysics training grant (T32 GM008294) and by the NSF GRFP. N.B. was supported in part by the NSF GRFP. J.D.Y. was supported by the Ruth L. Kirschstein National Research Service Award Postdoctoral Fellowships GM112294.

